# Structural damping renders the insect exoskeleton mechanically insensitive to non-sinusoidal deformations

**DOI:** 10.1101/2022.12.27.522009

**Authors:** Ethan S. Wold, James Lynch, Nick Gravish, Simon Sponberg

## Abstract

Muscles act through elastic and dissipative elements to mediate movement, but these elements can introduce dissipation and filtering which are important for energetics and control. The high power requirements of flapping flight can be reduced by the insect’s exoskeleton, which acts as a structurally damped spring under purely sinusoidal deformation. However, this purely sinusoidal dynamic regime does not encompass the asymmetric wing strokes of many insects or non-periodic deformations induced by external perturbations. As such, it remains unknown whether a structural damping model applies broadly and what implications it has for control. We used a vibration testing system to measure the mechanical properties of isolated *Manduca sexta* thoraces under symmetric, asymmetric, and band-limited white noise deformations. We measured a thoracic stiffness of 2980 *N*m^−1^ at 25 Hz and physiological peak-to-peak amplitude of 0.92 mm. Power savings and dissipation were indistinguishable between symmetric and asymmetric conditions, demonstrating that no additional energy is required to deform the thorax non-sinusoidally. Under white noise conditions, stiffness and damping were invariant with frequency, which is consistent with a structural damping model and suggests the thorax has no frequency-dependent filtering properties. A simple flat frequency response function fits our measured frequency response. This work demonstrates the potential of structurally damped materials to simplify motor control by eliminating any velocity-dependent filtering that viscoelastic elements usually impose between muscle and appendage.

## 1 Introduction

Muscles often act through series or parallel elastic elements to mediate movement, which have important consequences for locomotion. Locomotor springs such as tendons, ligaments, and exoskeleton serve many context-dependent energetic roles, including absorbing, recycling, and amplifying mechanical power (*1–3*). In fulfilling these roles, elastic elements and their associated structures in an organism can act as a mechanical filter between muscle and output motion, as well as between external stimuli and sensory feedback systems (*4–6*). The biomaterials that comprise musculoskeletal systems are also often significantly dissipative, introducing energy losses and phase offsets between actuator force and appendage motion (*7–9*). Nonideal transmissions such as latches and joints within locomotor systems can introduce additional energetic losses and contribute to local control of rapid movements by slowing the flow of quickly-released elastic energy (*10*, *11*).

To account for this biomechanical filtering, an animal must tune the muscle force magnitude and timing it uses to produce a given movement or respond to a perturbation. During steady, sinusoidal, and symmetric appendage movement, constant force and timing adjustments are largely unnecessary due to the single-frequency nature of these movements. However, steady locomotion often has asymmetric appendage movement (*12–14*). Furthermore, during unsteady locomotion, musculoskeletal systems must contend with dynamic loading that may contain a wide frequency band, impose transient perturbations and often require rapid, adaptive control (*15*). In these situations, if body mechanics operate in a significantly frequency-dependent way, it is likely that the neuromuscular system does as well.

The insect flight system presents a key example of how a highly complex elastic structure can affect energy flow during periodic movement. Many insects actuate their wings indirectly via deformations of the thoracic exoskeleton, which are created by the contraction of the flight power muscles (*16*). Spring-like properties of the exoskeleton can reduce the power requirements for flapping flight via elastic energy exchange if inertial power requirements are significant (*17–19*). While the thorax has been studied in this energetic context, the implications of thorax mechanics for control have been explored in far less detail (*20–22*).

Recent work has characterized the mechanical response of the hawkmoth *Manduca sexta* exoskeleton under purely sinusoidal deformations, showing that the exoskeleton is a linear spring with frequency-independent structural damping (*18*). Dissipation due to structural damping from cycle to cycle depends on the amplitude of oscillation and not the frequency. This contrasts with viscous damping which is a common model in biomaterials, especially muscle. During sinusoidal deformations, viscous materials dissipate energy per cycle proportional to both amplitude and frequency (*23*). Structural damping is thought to originate from internal friction between fibrous or filamentous structures, making it similar to Coulomb sliding friction which also dissipates energy per cycle dependent on amplitude, but not velocity (*24*). However, first-principles models of structural damping do not exist and most successful modeling attempts rely on the use of a phenomenological characterization, or restrictions to single-frequency motion (*25*).

While structural damping is present at the whole thorax level, this property has also been demonstrated in resilin, the constituent elastic protein that gives insect joints much of their resilience, as well as the whole joints of cockroaches during out-of-plane rotation (*26*, *27*). Recent modeling work has shown that structural damping is a key determinant of the dynamic efficiency of flapping insects (*28*). Altogether, while the presence of structural damping has been demonstrated in multiple insects, its relevance beyond steady, periodic conditions and impacts on motor control have not been explored. It is possible that viscous type damping emerges in non-sinusoidal regimes, which would give a drastically different prediction for how energy flows through the flight system. If the insect thorax is viscously damped under non-sinusoidal deformation, within-wingstroke frequency modulation would incur additional energetic cost, and require frequency-dependent tuning of muscle timing to compensate for phase lags. If a structural damping model holds in non-sinusoidal regimes, frequency modulation could be achieved with the same muscle force magnitude and phasing as in symmetric wingstrokes.

To investigate effects of non-sinusoidal deformations on thorax mechanics, we subjected *Manduca* thoraces to two different types of mechanical stimuli: asymmetric, periodic movements and band-limited white noise. We generated asymmetric displacement signals that create asymmetry between upstroke and downstroke, mimicking asymmetries observed in free flight (*29*). Then, we subjected the thorax to band-limited white noise to determine its response to displacement signals with a wide frequency content. This condition simulates potential wingstroke modulation that a hawkmoth may perform to remain stable in the air in response to aerodynamic perturbations, turbulence, or changing wind speeds (*29–31*). Similar wingstroke irregularities also may be introduced or exacerbated when the wings become damaged (*32–34*). We predict that the storage and dissipative characteristics of the thoracic spring will be the same for a given deformation amplitude, regardless of asymmetry or the frequency content of the deformation.

## 2 Materials and Methods

### Animals

*Manduca sexta* were obtained as pupae from colonies based at the University of Washington and Case Western Reserve University. Moths were stored in a humidity and temperature-controlled incubator on a 12 L: 12 D cycle. All animals were used within seven days of eclosion, and moth mass was equal to 2.24 ± 0.49 g. Both male and female moths were used.

### Thorax preparation

Our preparation was based on that of Gau, et al. 2019. Moths were anesthetized in a refrigerator for approximately 20 minutes, and the thorax was immediately isolated from the head and abdomen. The scales, wings, legs, and first thoracic segment were removed to isolate the second and third thoracic segments, which are the regions responsible for indirect actuation of the wings. The thoracic ganglion was carefully severed, accessed via the opening created by removing the head, to eliminate residual nervous system activity. Special care was taken to sever the flight muscle so that any spontaneous activity would not transmit appreciable force to the exoskeleton.

### Shaker mounting and preparation

We again follow the methods of Gau, et al. 2019 with minor adjustments. Custom 3D printed mounts were secured to the posterior phragma and the scutum with cyanoacrylate glue. The scutum mount was rigidly attached to a 10 N force transducer with a resonant frequency of 300 Hz (FORT1000, World Precision Instruments, Sarasota FL), and the posterior phragma mount was rigidly attached to the shaker head. The force transducer was attached to a micromanipulator, giving us precise control over the thorax’s rest length. Importantly, we aligned the thorax such that deformations occurred along the axis that the downstroke muscle would contract in the live animal (Fig. 1a). We measured the weight of the posterior mount and glue before securing it to the thorax and returned the system to this weight after mounting the thorax to the shaker. We then pre-compressed the thorax by 0.21 mm using the micromanipulator to match the average measured *in vivo* operating length of the DLM during tethered flight (*37*). An analog Hall effect sensor (DRV5053-Q1, Texas Instruments, Dallas TX) was used to measure displacement from a permanent magnet embedded in the shaker head. The Hall effect sensor was calibrated daily using a fifth order polynomial immediately prior to mounting the first specimen. The shaker head itself was driven by an electrodynamic vibration testing system (VTS600, Vibration Testing Systems, Aurora OH) (*36*).

**Figure 1:**
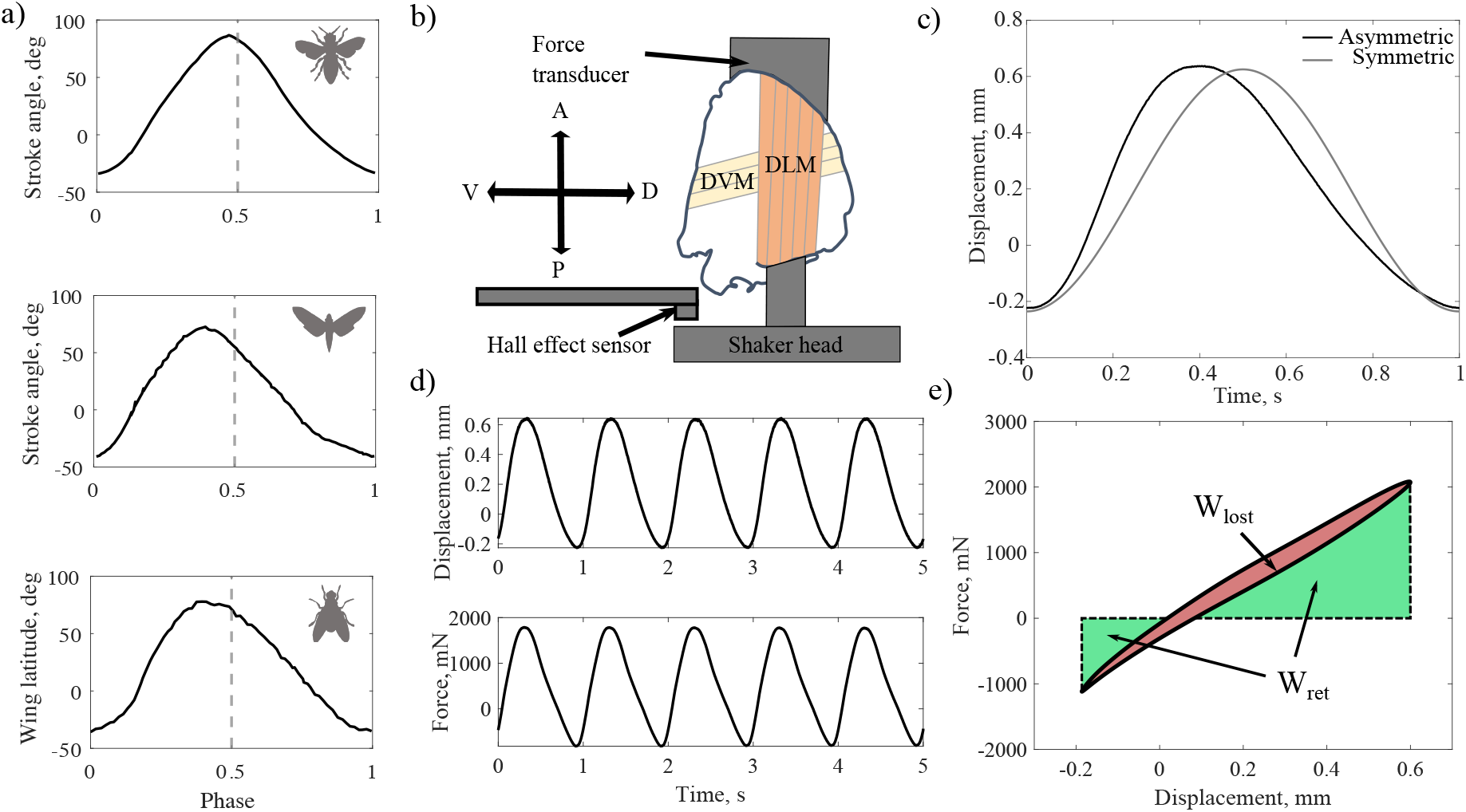
(a). Kinematic data from bumblebee (top), hawkmoth (middle), and fly (bottom) flight showing various degrees of non-sinusoidality and asymmetry (*14*, *29*, *35*). (b). Schematic of hawkmoth thorax mounted to the head of an electrodynamic shaker and a force transducer (*18*,*36*). The dorsolongitudinal (DLM, downstroke) and dorsoventral (DVM, upstroke) muscles are labeled and act through the thoracic shell to produce wingstrokes. A hall effect sensor measures displacement from a magnet embedded in the shaker head. (c) Example input signals for the symmetric and asymmetric deformations at 1 Hz. The asymmetric signal shows a clear difference in the rise and fall time of the oscillation. (d) Representative timeseries for an an asymmetric experiment at 1 Hz showing that asymmetries in the displacement waveform are recapitulated in the force waveform. (e) Representative force-displacement plot with dissipated and returned work highlighted in red and green respectively. The plot is not symmetric about the origin because the thorax has been pre-compressed.

### Deformations

We first prescribed sinusoidal deformations at both 1 and 25 Hz and a physiological peak-to-peak amplitude of 0.92 mm, equivalent to 4.5% amplitude strain of a typical 10 mm long thorax. The 25 Hz condition matches *Manduca’s* real wing beat frequency (29). While there is likely some small variation in strain amplitudes between individuals, the protocol of using a mean strain across all individuals has been established in the literature (37,38). From our length measurements, we calculated that the amplitude of our deformations correspond to 4.5 ± 0.15% thorax length across all individuals. We then prescribed an asymmetric sinusoid generated by splicing together two sine waves to create an upstroke and downstroke of different frequencies (Fig. 1b-c). The intrinsic nonlinear dynamics of our shaker filtered these signals, smoothing out the connections between each sinusoid, but preserved substantial asymmetry which was sufficient to test our hypotheses. Throughout we report the resulting strains rather than the prescribed ones. We tested asymmetry at an average frequency of 1 Hz and 25 Hz, although the precise nature of the asymmetry differed between frequencies due to the intrinsic mechanics of the shaker. The average frequency is defined as the inverse of the duration of one asymmetric upstroke and downstroke. In a few individuals, we implemented the asymmetry on thorax compression rather than extension to test whether the timing of the asymmetry within the overall oscillation affected thoracic energetics. We found no difference between the timings, so we proceeded to collect the complete dataset with a faster compression phase. Finally, we pre-scribed band-limited white noise for five seconds, with an upper frequency limit of 25 Hz. The shaker also filtered these signals, but the resulting deformations had substantial power density across the 25 Hz band. At frequencies above 25 Hz, we found it difficult to generate reliable asymmetries and white-noise with our shaker at our desired amplitudes, so we chose the in vivo wing beat frequency as our upper frequency limit.

### Data Analysis

We calculated components of mass-specific mechanical power by integrating the force-displacement curves, multiplying the resulting energies by the frequency of oscillation, and dividing by body mass (*18*) (Fig. 1d). We define *P_return_* to be the mechanical power savings from elastic energy storage and *P_lost_* to be the oscillation-averaged dissipated power. *P_in_* is the total mechanical power required to deform the thorax and is the sum of *P_return_* and *P_lost_*. For asymmetric conditions, we used the average frequency as defined above, either 1 or 25 Hz.

We utilize the structurally damped spring-damper material model previously established for cockroach leg joints and hawkmoth thorax to compute thorax material properties (*18*, *27*). This model consists of a spring with stiffness *k* in parallel with a damper that is characterized by a structural damping parameter *γ*. There are two common mathematical formulations: one utilizes a complex spring stiffness equal to *k*(1 + *γj*), while the other treats the damper as an equivalent viscous damper with a coefficient 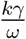 (*25*, *28*). These two models both result in frequency-independence and are equivalent for deformations of a single frequency, as we show below.

To calculate a spring stiffness and structural damping parameter, we applied a fast Fourier transform (FFT) to our force and displacement time series. We divided the transformed force in the frequency domain by the transformed displacement to calculate the complex modulus. The complex modulus is a frequency-dependent quantity, whose real part represents the elastic characteristics of a material, and imaginary part represents the dissipative characteristics. The effective stiffness (*k*) of the thorax was calculated from the real part of the complex modulus and the structural damping parameter, *γ* was calculated from the complex part, as shown in the following equations:

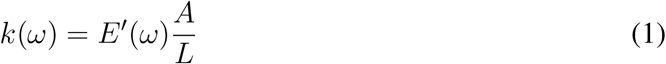

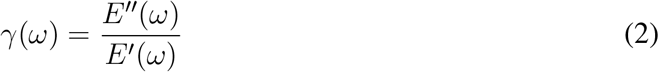

Here, *E′*(*ω*) is the real part of the complex modulus, *E″*(*ω*) is the imaginary part of the complex modulus, A is the area of muscle attachment, and L is thorax length. Since our experiments were carried out at either 1 Hz or 25 Hz, we extracted the material properties at these two frequencies.

For the white noise data, we applied a time-averaged FFT to force and displacement with an overlapping rectangular window. Like the asymmetry data, we compute a complex modulus by dividing the force and displacement in the frequency domain. Since the white noise stimulus contained frequencies up to 25 Hz, the complex modulus can be thought of as the frequency response of the thorax over this band. Using Newton’s second law for the vibrating thorax, we derived an analytical prediction for the frequency response function (FRF) of a structurally damped spring with negligible inertia,

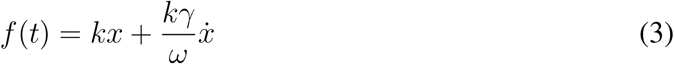

where *x* is the linear displacement of the thorax, *k* is the linear thorax stiffness, *γ* is the damping parameter, and *f*(*t*) is the force produced by deforming the thorax. We utilize the equivalent viscous damper formulation of a structural damper, which manifests in frequency-independent damping for sinusoidal motion. The velocity of a single-frequency sinusoidal trajectory (*ẋ*) will be linearly proportional to the frequency of oscillation. This factor of frequency cancels out with the 1/*ω* in the damping coefficient to yield a frequency-independent FRF. Taking the Laplace transform of both sides and solving for the frequency response function yields:

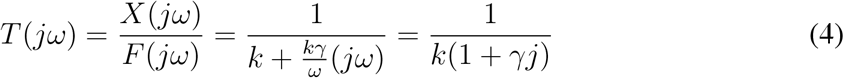

This frequency response function has a magnitude and phase given by the following:

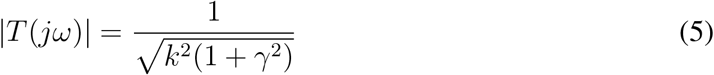

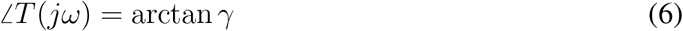

Under the assumption that *γ* << 1, these expressions predict a constant gain and phase of 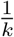 and *γ* respectively. Note that the frequency variable *ω* disappears from the magnitude and phase of the FRF, reflective of the frequency-independent property of a structural damper. To compare the energetics under asymmetric and symmetric conditions, we applied unpaired two-sample t-tests using a significance threshold of 0.05. All variation reported are one standard deviation from the mean.

## 3 Results

### 3.1 Symmetric and asymmetric deformations do not affect thorax material properties

Under symmetric deformation at 1 and 25 Hz (Fig. 2a,b) as well as asymmetric deformation at 1 Hz (Fig. 2c), thoraces exhibit characteristics of a weakly damped, linear spring as evidenced by the relatively constant slope of each loop. In the asymmetric 25 Hz case (Fig. 2d), there appears to be a slight downward curvature of the loop in tension compared to the symmetric case. Our sign convention is such that positive displacements represent compression while negative displacements represent tension.

**Figure 2:**
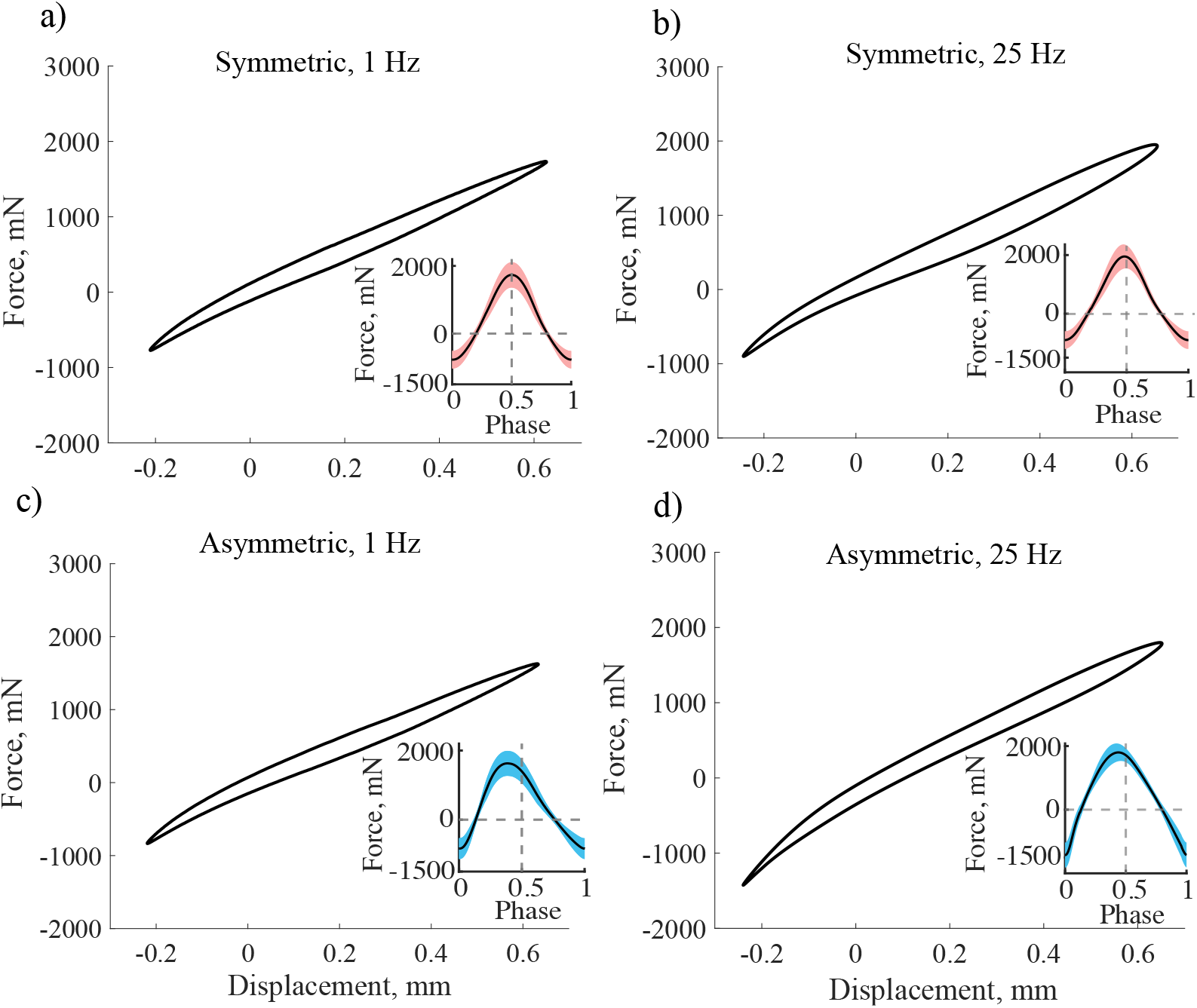
Force-displacement data across all experimental conditions. Mean forcedisplacement loops for all individuals under symmetric displacements at (a) 1 Hz (N=14) (b) and 25 Hz (N=14), and asymmetric displacements at (c) 1 Hz (N=14) (d) and 25 Hz (N=5) (d). Displacements across all conditions were 0.92 mm peak-to-peak. Oscillations at 25 Hz averaged slightly larger forces than oscillations at 1 Hz. Insets show mean ± standard deviation forces for a single oscillation.

To quantitatively compare the shapes of the symmetric and asymmetric force-displacement loops, we compute stiffness (1 Hz, sym: 2.93 ± 0.73 kNm^−1^; 1 Hz, asym: 3.05 ± 0.73 kNm^−1^; 25 Hz, sym: 2.78 ± 0.69 kNm^−1^; 25 Hz, asym: 3.33±0.68 kNm^−1^) and the structural damping factor (1 Hz, sym: 0.113±0.027; 1 Hz, asym: 0.118 ± 0.034; 25 Hz, sym: 0.095 ± 0.035; 25 Hz, asym: 0.101 ± 0.022) under each symmetry and frequency condition. We found no statistically significant differences between stiffness or damping at either 1 Hz (Fig. 3a) or 25 Hz (Fig. 3b), as determined by unpaired two-sample t-tests (stiffness: *p* = 0.66, *p* = 0.14; damping: *p* = 0.67, *p* = 0.71). The slight stiffening (10%) we observed in the 25 Hz asymmetric case compared to the symmetric case (Fig. 3b) may be the result of the nonlinear properties of the exoskeleton being excited by a higher frequency asymmetry, therefore resulting in the slight downward curvature of the force-displacement loop (Fig. 2d) and increased peak force during tension. All stiffness values are somewhat higher than the stiffness found by Gau *et al*, but result in peak forces within the range of the previously reported force-producing capacity for the *Manduca sexta* DLM (*38*).

**Figure 3:**
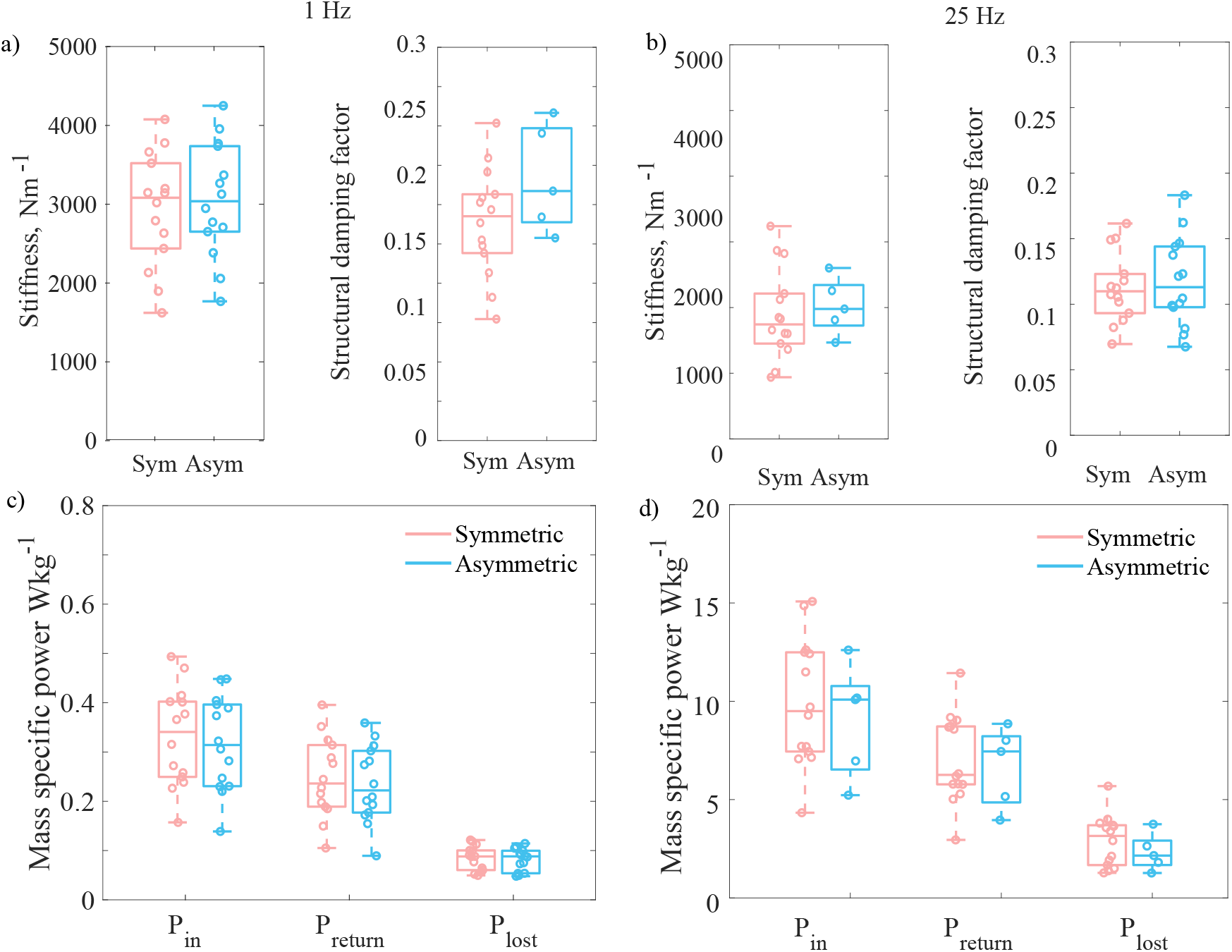
Material properties and thoracic powers for symmetric and asymmetric deformations. Stiffness and the structural damping constant do not differ between oscillation conditions at 1 Hz (a) and 25 Hz (b) regardless of whether the deformation is symmetric or asymmetric. Mass-specific total power expended, dissipated, and returned do not differ between symmetric and asymmetric conditions at either 1 Hz (c) or 25 Hz (d).

### 3.2 Cycle-averaged mechanical power is equivalent during symmetric and asymmetric deformations

To determine whether deviations from a sinusoidal trajectory modify thoracic energetics, we compared power returned (1 Hz, sym: 0.248 ± 0.082 Wkg^−1^; 1 Hz, asym: 0.235 ± 0.082 Wkg^−1^; 25 Hz, sym: 7.06 ± 2.25 Wkg^−1^; 25 Hz, asym: 6.69 ± 2.05 Wkg^−1^) and power lost (1 Hz, sym: 0.084 ± 0.025 Wkg^−1^; 1 Hz, asym: 0.082 ± 0.023 Wkg^−1^; 25 Hz, sym: 2.90 ± 1.29 Wkg^−1^; 25 Hz, asym: 2.32 ± 0.94 Wkg^−1^) during each symmetry and frequency condition. Returned and lost power were indistinguishable between symmetric and asymmetric conditions at both 1 Hz (*p* = 0.69, *p* = 0.20) and 25 Hz (*p* = 0.75, *p* = 0.38), which is consistent with our prediction that structural damping losses should only depend on deformation amplitude (Fig. 3d). The slight stiffening observed in Fig. 3b does not significantly affect power saved by the thoracic spring, therefore we suggest that it is not biologically relevant, at least under the conditions studied here.

### 3.3 The thorax has a flat frequency response curve

To assess whether a signal with a continuously changing frequency content modifies exoskeleton energetics, we subjected the thorax to band-limited white noise deformation signals (Fig. 4a). Filtering of our white noise signals by the shaker results in a spectrum that is not perfectly uniform over the band of interest, but still shows a clear cutoff frequency at 25 Hz (Fig. 4b). Regardless, our conclusions do not depend on the stimulus being perfectly white, since we are interested in testing the thorax’s response to a generalized irregular deformation.

**Figure 4:**
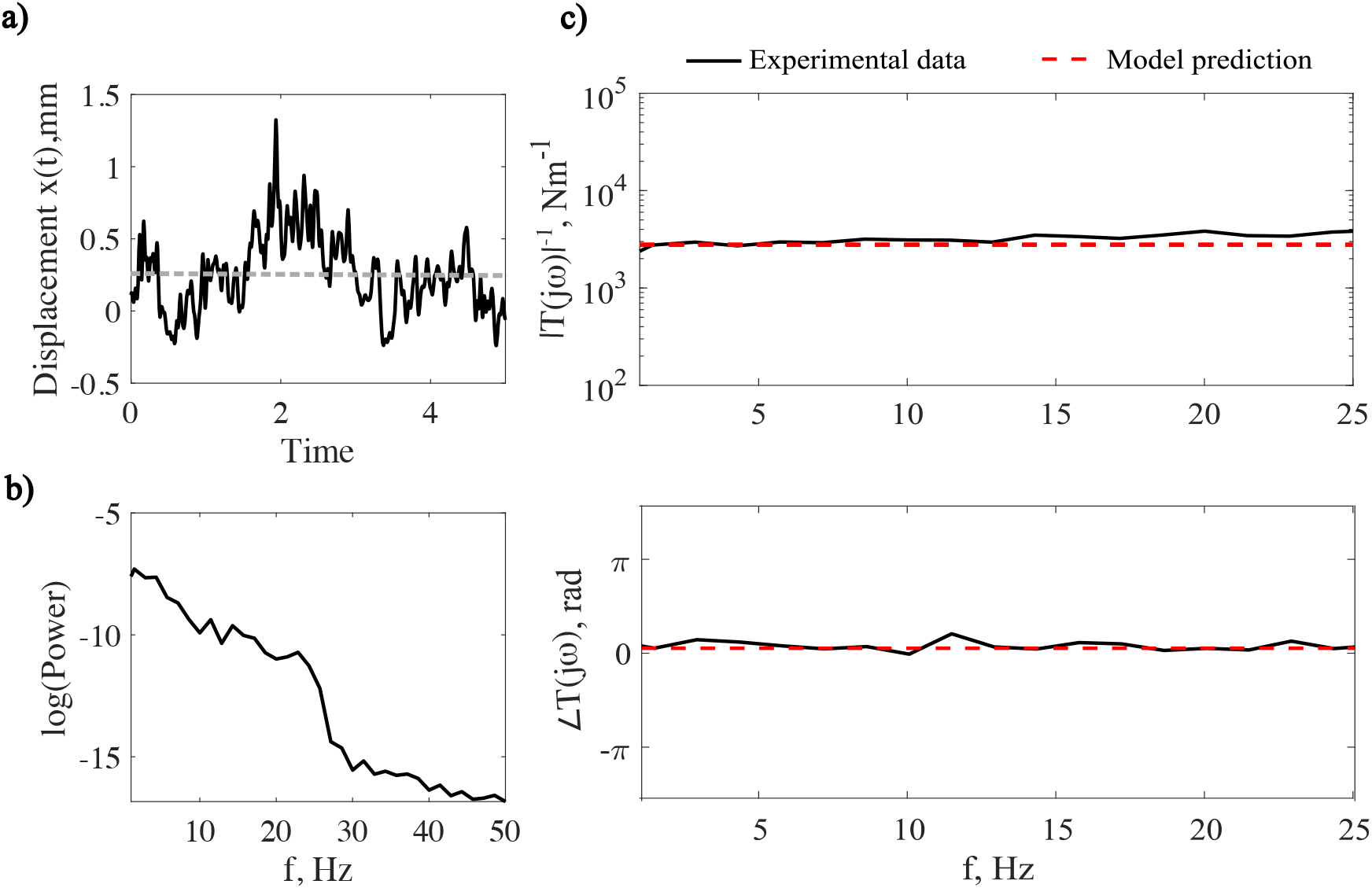
Frequency response of the thorax and characterization of damping. (a) A representative white noise displacement time series mimics an irregular perturbation with large frequency content. Note that the mean displacement shown by the grey dashed line is around 0.21 mm, corresponding to the prescribed thorax precompression. (b) The mean power spectrum of the detrended measured displacements in all individuals shows the majority of power within the 1-25 Hz band. (c) Gain and phase, which are analogous to stiffness and the damping constant, are flat with frequency, which is in accordance with the prediction from an analytical structurally damped frequency response function.

We measured the frequency response function in the thorax to have a gain and phase that are relatively flat with frequency (Fig. 4). We found an average gain and phase of 3.9 kNm^−1^ and 0.17 at 25 Hz, respectively. These values are in accordance with published stiffness and damping factors from hawkmoths and cockroaches (*18*, *27*). They are towards the higher end of the estimates of stiffness and damping from oscillatory deformation in this paper. Our theoretical prediction for a structurally damped spring with negligible inertia fits well on top of our measured data. Since our definitions of gain and phase correspond directly to stiffness and structural damping coefficient, these results are equivalent to stating that there is no substantial increase in stiffness or damping during irregular motion containing frequencies in the band we tested. However, data on symmetric oscillations of *Manduca* thoraces at frequencies up to 90 Hz lead us to believe that this trend likely extends significantly beyond 25 Hz limit imposed by our technological constraints (*18*).

## 4 Discussion

### 4.1 The thorax is mechanically insensitive to asymmetric oscillations

Our results demonstrate that there does not exist a significant additional energetic cost to deform the thorax asymmetrically. If velocity-dependent viscous damping was a dominant source of dissipation in the thorax, we would expect a distortion of the asymmetric force-displacement loop relative to the symmetric one that would manifest in a measurably greater lost power, since its frequency changes within a single period of oscillation. In contrast, we observe the same lost power as observed during symmetric deformations, as demonstrated by the same hysteretic loops (Fig. 2b, d). This energetic insensitivity to asymmetry persists at both physiologically relevant wingbeat frequencies for *Manduca* (25 Hz) and for frequencies an order of magnitude lower (1 Hz). One implication of these results is that observations of thorax mechanics at frequencies far below 25 Hz are likely representative of mechanics at 25 Hz. Measurements of insect thorax mechanics that can be made more easily at a low frequency may then be scaled up by frequency to draw conclusions about higher frequencies. While we did not explicitly test asymmetry at frequencies much higher than 25 Hz due to technical constraints, purely sinusoidal experiments have demonstrated frequency-independence up to at least 90 Hz in *Manduca* (*18*). Therefore, it is very likely that asymmetries are energetically inconsequential at higher frequencies as well.

Asymmetric upstroke and downstroke frequencies are a relatively common feature of insect flight and may serve important aerodynamic roles, despite not significantly affecting thoracic mechanics (*14*, *29*, *39*). Numerical modeling has demonstrated significant effects of upstroke-downstroke duration asymmetry on both thrust and lift production (*40*, *41*). In particular, faster upstrokes tend to increase thrust, which may explain the consistent observations of faster upstrokes in forward insect flight at varying speeds (*14*). Furthermore, the relative speeds of upstroke and downstroke have been linked to controlling body pitch (*42*). Therefore, a structurally damped thorax may not require additional muscle force tuning while adjusting duty factor to maneuver its body or generate enhanced forward propulsion. Our results suggest that insect flight models that approximate wingstrokes as symmetric likely generalize quite well at the level of thoracic mechanics, although this is likely not true for aerodynamics. Moreover while the mechanics remain the same, the effective inertia of the system may change with frequency because aerodynamic added mass scales with acceleration of the wing (*43*). Nonetheless the material properties are frequency independent.

### 4.2 Implications of a flat frequency response of the exoskeleton

Our white noise tests supports the hypothesis that the thorax acts like a structurally damped material even under irregular deformations that encompass a wide band of frequencies. The invariance of thorax stiffness and damping with frequency during white noise implies that there is no frequency dependent filtering of mechanical stimuli at least up to the natural wing beat frequency of 25 Hz, and likely higher given the properties of a structurally damped system. This result has implications for insects like *Manduca* that transiently modify their wingbeat frequency in the air on within-wingstroke timescales or experience transient perturbations.

Thoracic mechanics play an important role in dictating the resonant properties of a flapping insect. Substantial frequency modulation may indicate that an insect does not beat its wings at the resonant frequency of its flight system, since operation at resonance generally implies a large energetic cost for rapid frequency changes. Such off-resonance wingbeat frequencies may incur a trade-off between perturbation robustness and energetic efficiency (*28*, *44*). In synchronous insects like hawkmoths, a neural driving frequency determines wingbeat frequency and must be modified to facilitate wingbeat frequency changes. Therefore, significant frequency-dependent thorax mechanics may have to be accounted for by the nervous system to create a transient deviation from steady-state. For example, frequency-dependent stiffening would necessitate additional muscle force to drive the system more quickly. Frequency-dependent damping would necessitate force being applied at a different phase of activation than the normal phase at steady-state. For a hawkmoth, our results suggest that the same phase and magnitude of muscle force could achieve frequency modulation.

Recent theoretical advances in nonlinear elasticity have demonstrated that insects with substantial series compliance in their flight systems may exhibit wide, band-type resonance (*45*,*46*). The likely source of such series elastic effects is the wing hinge, which is also the primary dissipative component of the hawkmoth thorax (*18*). In these cases, there may be a range of frequencies over which efficient flight is possible, and frequency modulation within this range would not incur the loss of efficiency as a classical resonator. Estimates of frequency modulation in free-flying hawkmoths yield approximately ±15% of normal wingbeat frequency (*30*). A large degree of series compliance in a dissipative wing hinge, combined with substantial frequency modulation presents an extremely difficult control task for a viscously damped system, since the phase lag introduced by series elasticity would change with frequency. In contrast, a structurally damped wing hinge would mitigate some of this control difficulty by maintaining a constant phase lag. While frequency response measurements of the hawkmoth wing hinge do not exist, there is evidence of the structurally damped protein resilin in the hinges of hawk-moths, dragonflies, and locusts (*19*,*47*). Therefore, in broad categories of insects, a structurally damped exoskeleton may ease frequency-dependent energetic and control demands during frequency modulation.

In addition to frequency modulation, a frequency-independent thorax has ramifications for perturbation rejection and recovery. Insect wings collide with objects ranging from raindrops to plant stems during flight (*48*, *49*). Such perturbations to steady-state flight must be responded to by the flight muscles to maintain stability. The mechanical transparency (a flat frequency response) we observe between the wing and thorax in hawkmoths implies that a control force applied by the muscle to the thorax in response to a perturbation will be transmitted to the wing hinge after the same phase lag and with the same scaling, regardless of the speed of force application. Thus, modulation of muscle force would act on the wings in the same way, independent of frequency, potentially simplifying the control challenge that moth faces in matching neural control to motor output. This may be relevant for both perturbation rejection as well as other control challenges such as during turning maneuvers or when wings become damaged (*32*, *42*, *50*, *51*).

The flat frequency response of the thorax also has implications for the biomechanical pressures and constraints that apply over evolutionary timescales. Even within closely related subgroups of insects, there is sometimes wide variability in wingbeat frequency, such as in the Bombycoid moths, whose wingbeat frequency range spans nearly an order of magnitude from 8-64 Hz (*52*). A structurally damped thorax may reduce the amount of concomitant neuromuscular and exoskeletal coevolution necessary to evolve different wingbeat frequencies, contributing to the lability of wingbeat frequency as a functional trait. Furthermore, the evolution of an indirect flight system has likely contributed to the diversification of higher wingbeat frequency insects (*53*). An indirect flight system is the most common configuration for flight muscles in insects, with the only significant exception being the Odonates (*53*). Previous proposed advantages to the indirect configuration involve the ability to actuate large amplitude wing strokes with extremely small muscle strains, making use of complex three-dimensional shape changes to create additional mechanical advantage (*21*,*54*). If the thorax introduced substantial velocity-dependent filtering and dissipation, this would present an obvious detriment to the indirect flight configuration that would require additional coevolution from the flight muscles and neural circuitry to circumvent. Thus, it is reasonable that the presence of structural damping in the exoskeleton of primitive insects may have contributed to the ubiquity of indirect flight as a preferred method of wing actuation in extant insects.

### 4.3 Implications of structural damping for non-flapping movement

Frequency-independent damping likely has relevance in organisms that do not utilize flapping aerial locomotion (*55*). There appear to be significant dynamic regimes in which vertebrate connective tissues such as tendons are largely frequency-independent. Such frequency independence has been observed at higher strain rates, while classic viscous behavior becomes more important at lower strain rates (*56*, *57*). The origin of this mechanical behavior remains unclear, however it may be related to sliding between collagen fibers or interactions between collagen and proteoglycan matrix (*58*). Frequency independence, potentially from structural damping, in these materials may be critical for high-power or impulsive movements in which viscous effects would either dissipate impractically large or dangerous amounts of energy, or interfere with control of rapid motion.

Despite having some functional similarity, the muscle architecture and elastic elements present in vertebrate locomotor systems differ in a few key ways from those in insects. Many insect muscles, such as the indirect flight muscles in hawkmoths, attach directly to the exoskeleton without any coupling elastic element (*53*). In vertebrates, series elastic tendons introduce velocity-dependent dynamics that can significantly muscle-tendon unit behavior, decoupling muscle and muscle-tendon unit dynamics. (*8*, *59*, *60*). In insects, analogous structures known as apodemes exist, but are over an order of magnitude stiffer on average than tendons. That vertebrate connective tissues are materially distinct from chitinous exoskeleton, but exhibit functionally similar characteristics under high-frequency loading illustrates an interesting commonality; however, it remains unclear what drives the emergence of this property across disparate species.

Fast, periodic movements encompass a large diversity of organismal locomotion (such as flapping flight), but differ fundamentally from ballistic movements performed by many animals. Recent work on ballistic locomotion has sparked a renewed usage of spring-mass-damper models to predict the flow of power from actuator input to motor output through the LaMSA framework (*10*, *61*, *62*). In LaMSA systems, power amplification is achieved by slowly transferring muscle contractile energy to a spring over a long time scale and releasing it extremely quickly to surpass inherent force-velocity muscle power limitations. Many of these models ignore internal damping altogether, or only consider velocity-dependent effects (*10*, *63*). In such “superfast” movements, consideration of different models of damping may give rise to altered dynamics, given the high velocities and lack of explicit muscular control after the initial loading of the spring. A structurally damped ballistic limb movement would likely dissipate much less energy than an equivalent viscously damped one, resulting in more energy delivered to the desired output, or unwanted internal damage. Many of these movements are generated by invertebrates that utilize bending of stiff exoskeleton as the LaMSA spring, such as the mantis shrimp raptorial appendage, and therefore may have the necessary microstructural properties that give rise to structural damping (*64*,*65*). Our work demonstrates a need to investigate multiple internal damping models in other systems that rely on complex energy flow to produce movement.

### 4.4 Conclusion

We find the mechanical response of the hawkmoth thorax in non-sinusoidal regimes to be consistent with phenomenological and mathematical predictions from a simple structural damping model, providing evidence that structural damping is a robust property of the thorax that extends beyond symmetric periodic conditions. The frequency-independent characteristics of structural damping render the thorax insensitive to non-sinusoidality and dynamically transparent to force inputs, allowing energy to flow directly between muscle and wing, despite the large elastic structure that facilitates indirect flight actuation.

## Acknowledgments

We thank Scott Wilburn, Dr. Paul Umbanhowar and Dr. Dan Goldman for access to the shaker apparatus and assistance with troubleshooting. This work was supported by US National Science Foundation RAISE grant no. IOS-2100858 to S.S. and N.G. and 1554790 (MPS-PoLS) and a Dunn Family Professorship to S.S. as well as the US National Science Foundation Physics of Living Systems SAVI student research network (GT node grant no. 1205878). Raw data are available from the Dryad Digital Repository at:

## Author Contributions

E.S.W., J.L., N.G., S.S. conceived and designed the study and edited the manuscript. E.S.W. ran experiments, analyzed data, and prepared the figures. All authors wrote and edited the manuscript.

## References

1. Roberts TJ, Azizi E. 2011 Flexible mechanisms: The diverse roles of biological springs in vertebrate movement. Journal of Experimental Biology 214, 353–361.

2. Griffiths RI. 1991 Shortening of muscle fibres during stretch of the active cat medial gastrocnemius muscle: the role of tendon compliance.. The Journal of Physiology 436, 219–236.

3. Konow N, Azizi E, Roberts TJ. 2012 Muscle power attenuation by tendon during energy dissipation. Proceedings of the Royal Society B: Biological Sciences 279, 1108–1113.

4. Tytell ED, Holmes P, Cohen AH. 2011 Spikes alone do not behavior make: Why neuroscience needs biomechanics. Current Opinion in Neurobiology 21, 816–822.

5. Chiel HJ, Ting LH, Ekeberg Hartmann MJ. 2009 The brain in its body: Motor control and sensing in a biomechanical context. Journal of Neuroscience 29, 12807–12814.

6. Cowan NJ, Fortune ES. 2007 The critical role of locomotion mechanics in decoding sensory systems. Journal of Neuroscience 27, 1123–1128.

7. Zajac FE. 1989 Muscle and tendon: properties, models, scaling, and application to biomechanics and motor control..

8. Robertson BD, Sawicki GS. 2015 Unconstrained muscle-tendon workloops indicate resonance tuning as a mechanism for elastic limb behavior during terrestrial locomotion. Proceedings of the National Academy of Sciences of the United States of America 112, E5891–E5898.

9. Tytell ED, Carr JA, Danos N, Wagenbach C, Sullivan CM, Kiemel T, Cowan NJ, Ankarali MM. 2018 Body stiffness and damping depend sensitively on the timing of muscle activation in lampreys. Integrative and comparative biology 58, 860–873.

10. Ilton M, Saad Bhamla M, Ma X, Cox SM, Fitchett LL, Kim Y, Koh Js, Krishnamurthy D, Kuo CY, Temel FZ, Crosby AJ, Prakash M, Sutton GP, Wood RJ, Azizi E, Bergbreiter S, Patek SN. 2018 The principles of cascading power limits in small, fast biological and engineered systems. Science 360.

11. Divi S, Ma X, Ilton M, St. Pierre R, Eslami B, Patek SN, Bergbreiter S. 2020 Latch-based control of energy output in spring actuated systems. Journal of the Royal Society Interface 17.

12. Herrera-Amaya A, Seber EK, Murphy DW, Patry WL, Knowles TS, Bubel MM, Maas AE, Byron ML. 2021 Spatiotemporal Asymmetry in Metachronal Rowing at Intermediate Reynolds Numbers. Integrative and Comparative Biology 61, 1579–1593.

13. Cavagna GA. 2010 Symmetry and asymmetry in bouncing gaits. Symmetry 2, 1270–1321.

14. Ellington C. 1984 The aerodynamics of hovering insect flight III: kinematics. Philosophical Transactions of the Royal Society B: Biological Sciences 305, 41–78.

15. Biewener AA, Daley MA. 2007 Unsteady locomotion: Integrating muscle function with whole body dynamics and neuromuscular control. Journal of Experimental Biology 210, 2949–2960.

16. Pringle S. 1940 The excitation and contraction of the flight muscles of insects.. Journal of Physiology pp. 226–232.

17. Dickinson MH, Lehmann FO, Chan WP. 1998 The control of mechanical power in insect flight. American Zoologist 38, 718–728.

18. Gau J, Gravish N, Sponberg S. 2019 Indirect actuation reduces flight power requirements in Manduca sexta via elastic energy exchange. Journal of the Royal Society Interface 16.

19. Weis-Fogh T. 1960 A Rubber-Like Protein in Insect Cuticle. Journal of Experimental Biology 37, 889–907.

20. Ando N, Kanzaki R. 2016 Flexibility and control of thorax deformation during hawkmoth flight. Biology Letters 12.

21. Deora T, Gundiah N, Sane SP. 2017 Mechanics of the thorax in flies. Journal of Experimental Biology 220, 1382–1395.

22. Jankauski MA. 2020 Measuring the frequency response of the honeybee thorax. Bioinspiration and Biomimetics 15.

23. Weaver Jr W, Timoshenko SP, Young DH. 1991 Vibration problems in engineering. John Wiley & Sons.

24. Drucker DC. 2021 Coulomb Friction, Plasticity, and Limit Loads. Journal of Applied Mechanics 21, 71–74.

25. Muravskii GB. 2004 On frequency independent damping. Journal of Sound and Vibration 274, 653–668.

26. Gosline J, Lillie M, Carrington E, Guerette P, Ortlepp C, Savage K. 2002 Elastic proteins: Biological roles and mechanical properties. Philosophical Transactions of the Royal Society B: Biological Sciences 357, 121–132.

27. Dudek DM, Full RJ. 2006 Passive mechanical properties of legs from running insects. Journal of Experimental Biology 209, 1502–1515.

28. Lynch J, Gau J, Sponberg S, Gravish N. 2021 Dimensional analysis of spring-wing systems reveals performance metrics for resonant flapping-wing flight. Journal of the Royal Society Interface 18.

29. Willmott AP, Ellington CP. 1997 The mechanics of flight in the hawkmoth Manduca sexta I. Kinematics of hovering and forward flight. Journal of Experimental Biology 200, 2705–2722.

30. Gau J, Gemilere R, Lds-Vip, Lynch J, Gravish N, Sponberg S. 2021 Rapid frequency modulation in a resonant system: Aerial perturbation recovery in hawkmoths. Proceedings of the Royal Society B: Biological Sciences 288.

31. Ortega-Jimenez VM, Greeter JS, Mittal R, Hedrick TL. 2013 Hawkmoth flight stability in turbulent vortex streets. Journal of Experimental Biology 216, 4567–4579.

32. Fernández MJ, Springthorpe D, Hedrick TL. 2012 Neuromuscular and biomechanical compensation for wing asymmetry in insect hovering flight. Journal of Experimental Biology 215, 3631–3638.

33. Fernández MJ, Driver ME, Hedrick TL. 2017 Asymmetry costs: Effects of wing damage on hovering flight performance in the hawkmoth Manduca sexta. Journal of Experimental Biology 220, 3649–3656.

34. Kihlstrom K, Aiello B, Warrant E, Sponberg S, Stockl A. 2021 Wing damage affects flight kinematics but not flower tracking performance in hummingbird hawkmoths. Journal of Experimental Biology 224.

35. Zanker JM. 1990 The wing beat of Drosophila Melanogaster. I. Kinematics. Philosophical Transactions of the Royal Society of London. B, Biological Sciences 327, 1–18.

36. Gravish N, Franklin SV, Hu DL, Goldman DI. 2012 Entangled granular media. Physical Review Letters 108, 1–4.

37. Tu MS, Daniel TL. 2004a Cardiac-like behavior of an insect flight muscle. Journal of Experimental Biology 207, 2455–2464.

38. Tu MS, Daniel TL. 2004b Submaximal power output from the dorsolongitudinal flight muscles of the hawkmoth Manduca sexta. Journal of Experimental Biology 207, 4651–4662.

39. Weis-Fogh T, Jensen M. 1956 Biology and physics of locust flight I. Basic principles in insect flight. A critical review.. Philosophical Transactions of the Royal Society B: Biological Sciences.

40. Yu Y, Tong B. 2005 Flow control mechanism in wing flapping with stroke asymmetry during insect forward flight. Acta Mechanica Sinica/Lixue Xuebao 21, 218–227.

41. Younsi A, El-Hadj AA, Abd Rahim SZ, Rezoug T. 2022 Effects of the wing spacing, phase difference, and downstroke ratio on flapping tandem wings. Proceedings of the Institution of Mechanical Engineers, Part C: Journal of Mechanical Engineering Science 236, 4689–4712.

42. Cheng B, Deng X, Hedrick TL. 2011 The mechanics and control of pitching manoeuvres in a freely flying hawkmoth (Manduca sexta). Journal of Experimental Biology 214, 4092–4106.

43. Sane SP. 2003 The aerodynamics of insect flight. Journal of Experimental Biology 206, 4191–4208.

44. Gau J, Wold ES, Lynch J, Gravish N, Sponberg S, Sponberg S. 2022 The hawkmoth wing-beat is not at resonance. pp. 1–5.

45. Pons A, Beatus T. 2022a Distinct forms of resonant optimality within insect indirect flight motors. Journal of The Royal Society Interface 19.

46. Pons A, Beatus T. 2022b Elastic-bound conditions for energetically optimal elasticity and their implications for biomimetic propulsion systems. Nonlinear Dynamics 108, 2045–2074.

47. Hollenbeck AC, Palazotto AN. 2013 Mechanical Characterization of Flight Mechanism in the Hawkmoth Manduca Sexta. Experimental Mechanics 53, 1189–1199.

48. Dickerson AK, Shankles PG, Madhavan NM, Hu DL. 2012 Mosquitoes survive raindrop collisions by virtue of their low mass. Proceedings of the National Academy of Sciences of the United States of America 109, 9822–9827.

49. Foster DJ, Cartar RV. 2011 What causes wing wear in foraging bumble bees?. Journal of Experimental Biology 214, 1896–1901.

50. Mountcastle AM, Alexander TM, Switzer CM, Combes SA. 2016 Wing wear reduces bumblebee flight performance in a dynamic obstacle course. Biology Letters 12, 1–4.

51. Beatus T, Guckenheimer JM, Cohen I. 2015 Controlling roll perturbations in fruit flies. Journal of the Royal Society Interface 12.

52. Aiello BR, Sikandar UB, Minoguchi H, Bhinderwala B, Hamilton CA, Kawahara AY, Sponberg S. 2021 The evolution of two distinct strategies of moth flight. Journal of the Royal Society Interface 18.

53. Dudley R. 2002 The biomechanics of insect flight: form, function, evolution. Princeton University Press.

54. Boettiger E, Furshpan E. 1952 The mechanics of flight movements in Diptera. Biological Bulletin pp. 200–211.

55. Ker RF. 1977 Some structural and mechanical properties of locust and beetle cuticle. PhD thesis University of Oxford.

56. Rosario MV, Roberts TJ. 2020 Loading Rate Has Little Influence on Tendon Fascicle Mechanics. Frontiers in Physiology 11, 1–9.

57. Crisco JJ, Moore DC, Mcgovern RD. 2002 Strain-rate sensitivity of the rabbit MCL diminishes at traumatic.pdf. 35, 1379–1385.

58. Bonner TJ, Newell N, Karunaratne A, Pullen AD, Amis AA, M.J. Bull A, Masouros SD. 2015 Strain-rate sensitivity of the lateral collateral ligament of the knee. Journal of the Mechanical Behavior of Biomedical Materials 41, 261–270.

59. Duenwald SE, Vanderby R, Lakes RS. 2009 Viscoelastic relaxation and recovery of tendon. Annals of Biomedical Engineering 37, 1131–1140.

60. Daley MA, Biewener AA. 2011 Leg muscles that mediate stability: Mechanics and control of two distal extensor muscles during obstacle negotiation in the guinea fowl. Philosophical Transactions of the Royal Society B: Biological Sciences 366, 1580–1591.

61. Longo SJ, Cox SM, Azizi E, Ilton M, Olberding JP, St Pierre R, Patek SN. 2019 Beyond power amplification: Latch-mediated spring actuation is an emerging framework for the study of diverse elastic systems. Journal of Experimental Biology 222, 1–10.

62. Sutton GP, Mendoza E, Azizi E, Longo SJ, Olberding JP, Ilton M, Patek SN. 2019 Why do large animals never actuate their jumps with latch-mediated springs? because they can jump higher without them. Integrative and Comparative Biology 59, 1609–1618.

63. Bolmin O, Socha JJ, Alleyne M, Dunn AC, Fezzaa K, Wissa AA. 2021 Nonlinear elasticity and damping govern ultrafast dynamics in click beetles. Proceedings of the National Academy of Sciences of the United States of America 118.

64. Farley GM, Wise MJ, Harrison JS, Sutton GP, Kuo C, Patek SN. 2022 Erratum: Correction: Adhesive latching and legless leaping in small, worm-like insect larvae (The Journal of experimental biology (2019) 222 Pt 15 PII: jeb243841). The Journal of experimental biology 225.

65. Patek SN, Korff WL, Caldwell RL. 2004 Deadly strike mechsanism of a mantis shrimp. Nature 428, 819–820.

